# MrHAMER2: high-accuracy long-read RNA sequencing to decode isoform-specific variation in viral transcripts during latency

**DOI:** 10.1101/2024.12.19.629526

**Authors:** Christian M. Gallardo, Jessica L. Albert, Andrew A. Qazi, Roni Lobato Ventura, Savitha Deshmukh, Nadejda Beliakova-Bethell, Bruce E. Torbett

## Abstract

Alternative splicing (AS) greatly expands the repertoire of proteins encoded by the human genome. Viruses have been shown to hijack AS cellular pathways to sustain replication or lead to latency. In HIV-1 infection, the virus integrates into the host genome, becoming a transcriptional unit that directly engages in AS to regulate its gene expression. Sequencing advances have enabled insights into HIV-1 gene expression dynamics during productive replication. However, viral isoform dynamics during latency remain largely uncharacterized due to the low abundance of spliced viral transcripts in associated CD4+ T cell subsets, making their accurate detection and quantification challenging. MrHAMER2 is a high-accuracy long-read RNA sequencing method that leverages dual Unique Molecular Identifier (UMI) tagging of cDNA to accurately capture and quantify full-length isoforms with high dynamic range and 99.968% single-nucleotide accuracy. We used MrHAMER2 to decode the spliced HIV-1 transcriptome in a primary CD4+ T cell model of latency and showed substantial changes in viral isoforms bearing intron retentions accompanied by changes in their potential to generate translatable protein.

## INTRODUCTION

Alternative splicing (AS) is a post-transcriptional regulatory mechanism that expands the repertoire of proteins encoded by the human genome^1^. During AS, exons from the same transcriptional unit are joined in different combinations to yield mRNA transcripts with similar, dissimilar, or at times mutually exclusive functions^2^. Given the importance of AS in regulating human gene expression, it is not surprising that a number of viruses hijack this process to promote their replication^3^ or dormancy.

During HIV-1 infection, the virus integrates into the host genome, thereby becoming a transcriptional unit that actively engages in AS to regulate viral gene expression. During this process, the virus co-opts host spliceosomal components so that a single 9.2 kb viral RNA (which codes for both the viral genome and Gag and Gag-Pol polyproteins) is alternatively spliced to place the open reading frames (ORFs) of distinct viral genes in close proximity to the 5’ cap, thus coding for the remaining viral proteins^4^. This atypical mode of gene expression regulation has additional features that exacerbate its splicing complexity compared to human genes, including overlapping ORFs, high dynamic range of viral transcripts, shared UTRs, and low transcript enrichment compared to host transcripts^5, 6^. Thus, viral gene expression results in prodigious viral isoform diversity with a large dynamic range of expression, resulting in up to 100 isoforms from a single transcriptional unit^7^. Due to the ensuing complex viral splicing landscape, long-read sequencing is invariably required to unambiguously decode the full breadth of viral isoform sequence variation^7-9^.

Advances in next-generation sequencing have brought about greater insights into how HIV-1 regulates its gene expression during active viral replication^10, 11^, with recent studies using emergent long-read sequencing approaches further closing knowledge gaps^5, 9, 12^. However, viral isoform regulation during HIV-1 latency remains largely uncharacterized despite major strides in the understanding of associated host cell transcriptional programs^13^. This knowledge gap results from the complex splicing of viral RNA and is exacerbated by the marked reductions in both spliced viral mRNA amounts and their originating CD4+ T cell subsets when infected cells transition to a latent state^13^. This makes the accurate detection and quantification of target viral isoforms from these cellular extracts exceptionally challenging during latency.

We developed MrHAMER2, a high-accuracy long-read RNA sequencing (RNA-seq) method to quantitatively capture full-length spliced isoforms from biologically relevant samples containing a low abundance of target transcripts within scarce CD4+ T cell subsets. The MrHAMER2 protocol is based on the long-read RNA sequencing foundations that were previously developed by us, including the previous generation of this protocol^14^ (Multi-read Hairpin Mediated Error-correction Reaction), optimized reverse transcription (RT) and emulsion PCR conditions^14^, and long cDNA enrichment capabilities via chemical ablation of 3’ RNA ends^5^. These existing methods were combined with the use of dual UMI tagging of single cDNA molecules, which has not previously been adopted in the context of long-read RNA-sequencing, to enable accurate amplification of rare isoforms from complex cellular transcript extracts while controlling for PCR sampling and chimerism artefacts. We show that MrHAMER2 can accurately detect and quantify full-length isoforms with high dynamic range and 99.968% single-nucleotide accuracy (Q35 equivalent). We extensively validated isoform and splice junction quantification and long-range phasing capabilities by benchmarking against PCR-free reference datasets and by using ad-hoc mixtures of synthetic RNA transcripts containing long-range mutation pairs. We then used MrHAMER2 to decode the spliced HIV-1 transcriptome in a primary CD4+ T cell derived viral latency model and in autologous T cell subsets. We show substantial changes in viral isoforms bearing intron retentions (IRs) in both latently-infected cells and in CD4+ T cell subsets sorted from this bulk population. We further demonstrate isoform productivity differentials that could severely affect the translatability and ultimately the function of certain viral proteins in CD4+ T cells.

## RESULTS

### Optimizing Dual UMI tagging to enable single-molecule long-read RNA sequencing

Dual UMI-based approaches have not been previously applied to long-read RNA-seq, particularly in the context of nanopore sequencing, so determining UMI tagging parameters was necessary to ensure the precise amplification of the spliced HIV-1 transcriptome from cellular extracts. Since each UMI tag contains a synthetic priming site, we used PCR amplification as a readout for UMI incorporation. We first tried adding the 3’ UMI via a flanking region in the Oligo-d(T) primer during reverse transcription (RT), followed by 5’ UMI addition via gene-specific priming (GSP) during second-strand synthesis. Even though this approach yielded UMI-tagged cDNA amplicons of the expected size when using RNA from productively-infected cells **(Supplementary Fig. 1A)**, the approach showed low specificity when using RNA from resting latently-infected cells that are the focus of our studies **(Supplementary Fig. 1B)**. We reasoned that coupling the 3’ UMI tagging with Oligo-d(T) priming was generating interference from the more highly enriched host cell mRNAs present in the total RNA extracts. Next, we attempted to improve 3’ UMI tagging specificity by using a tailed GSP approach during RT, but the only approach that yielded improvements was running a conventional Oligo-d(T) primed RT, followed by sequential addition of 5’/3’ UMI tags via GSP during second strand synthesis **(Supplementary Fig. 1C)**.

We integrated the UMI-tagging optimizations into a finalized assay scheme (**Fig. 1A**). In the optimized workflow, total RNA is treated with the CASPR (Chemical Ablation of Spuriously Priming RNAs) reagent^5^, followed by Oligo-d(T_20_) primed RT using SuperScript IV. Both 5’ and 3’ UMI tags, each containing synthetic priming sites, are added during second-strand synthesis by targeting the 5’/3’ UTR regions (shared in all HIV transcripts) with GSP. Dual UMI-tagged cDNA molecules are then amplified via emulsion PCR (emPCR) targeting the added synthetic priming sites flanking the UMIs. Libraries of the resulting amplicons are then prepared using the Oxford Nanopore Technologies (ONT) Native Barcoding kit and sequenced using the new Q20+ nanopore chemistries. The resulting reads are processed using our bioinformatic pipeline **(Fig. 1B)**, which extracts the UMIs and clusters associated reads based on their UMI identity. The resulting UMI clusters, each originating from a single Dual UMI-tagged cDNA molecule, are filtered to have a minimum cluster size, assembled, and then polished to yield high accuracy single molecule sequences (referred to as MrHAMER2 reads). This bioinformatic pipeline is coupled to a post-hoc chimera filter module **(Supplementary Fig. 2)**, conceptually modeled after Karst et al. 2022^15^ that leverages the dual UMIs to identify single molecule clusters that arose due to PCR recombination, a phenomenon that we and others have previously shown to be a pervasive source of sequence artefacts^14, 16^. Unlike other long-read UMI-based pipelines, MrHAMER2 yields FASTQ files that are amenable to sequence processing and filtering in various downstream applications.

**Figure 1.**
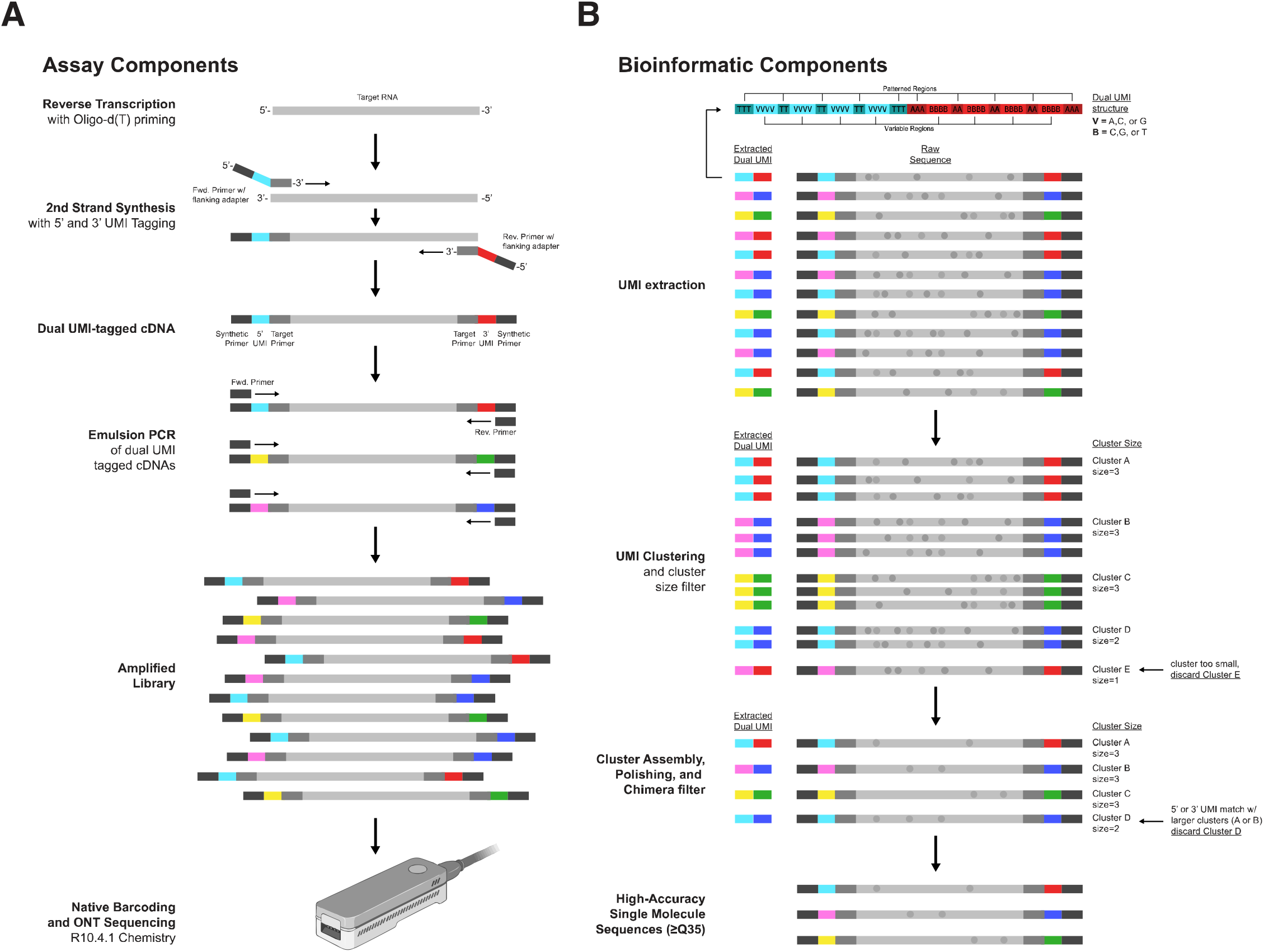
MrHAMER2 assay and bioinformatic components. **(A)** The assay component first involves reverse transcription of target RNAs with Oligo-d(T) priming. UMIs containing flanking synthetic priming sites are added to both 5’ and 3’ ends of the first-strand cDNA via gene-specific priming using respective 5’/3’ UTR sequences shared in all target isoforms. Dual UMI-tagged molecules are amplified via emulsion PCR, and libraries are prepared from the resulting amplicons using the Native Barcoding kits from ONT followed by Nanopore sequencing using R.10.4.1 chemistry. **(B)** The bioinformatic component involves extraction of the patterned UMIs from each read, followed by clustering of reads into bins based on dual UMI identity. A minimum cluster size filter is then applied so that a minimum number of balanced (i.e. sense and antisense) raw reads are available per dual UMI cluster to ensure sufficient error-correction. Dual UMI clusters passing the size filter are then assembled and polished into high-accuracy consensus assemblies originating from single cDNA molecules. A post-hoc chimera filtering module is then used to identify partial UMI matches present in other clusters and remove the less enriched recombinant clusters, resulting in high-accuracy single molecule sequences free of PCR sampling or chimeric artefacts.

### MrHAMER2 yields highly accurate single-molecule long-read RNA sequences

We evaluated the accuracy improvements elicited by the MrHAMER2 pipeline by using long *in vitro* transcribed (IVT) RNA of known sequence. As expected, we found that cluster size (i.e. the number of passes per single molecule) is directly proportional to single molecule accuracy as measured by Q-score, but also inversely proportional to MrHAMER2 reads yield **(Fig. 2A)**. We find that a minimum cluster size of 4 balances accuracy and yield of MrHAMER2 reads, resulting in ∼Q35 accuracy. Accuracy tapered off at ∼Q38.5 at cluster size of ≥8 compared to the maximum Q42.5 found when using DNA inputs **(Supplementary Fig. 3A)**. This suggests the maximum theoretical error rate of MrHAMER2 when using RNA inputs is limited by the combined error rate of the RNA polymerase and reverse transcriptase^17^. When using a minimum cluster size of 4, MrHAMER2 reduces raw Nanopore error (when using the R.10.4.1 chemistry) by 46-fold, from 1.472% to 0.0320% **(Fig. 2B)**. We also tested dual UMI assisted duplex basecalling post-hoc and reached Q33 accuracy **(Supplementary Fig. 3B)** and higher MrHAMER2 read yield for applications that require more throughput. To test for long-range accuracy, we used a set of IVT RNA inputs containing a known 8 bp barcode at either the 5’ or 3’ end of the molecule and mixed them at 1:1 ratio, then sequenced with MrHAMER2 to count recombinant sequences containing both barcodes or no barcodes **(Supplementary Fig. 3C)**. We found that our chimera filtering scheme combined with the use of emPCR reduces template switching from a high of 56.59% in Aqueous PCR (aqPCR) with 200K RNA input molecules, to undetectable levels across a range of RNA inputs **(Fig. 2C)**. To validate ability to capture long-range linkage across a range of enrichment levels, three mutants were generated containing pairs of distant single-nucleotide variants (SNVs), mixed in known proportions, then used as inputs for MrHAMER2 sequencing **(Fig. 2D)**. The resulting haplotypes show that the observed frequencies of each mutant are comparable to expected frequencies for the range of RNA inputs from 2K to 200K, with R^2^ values of 0.99 for all treatments **(Fig. 2E)**. These results demonstrate that MrHAMER2 can accurately capture long RNA sequence variation across a range of concentrations and without detectable sequence artefacts.

**Figure 2.**
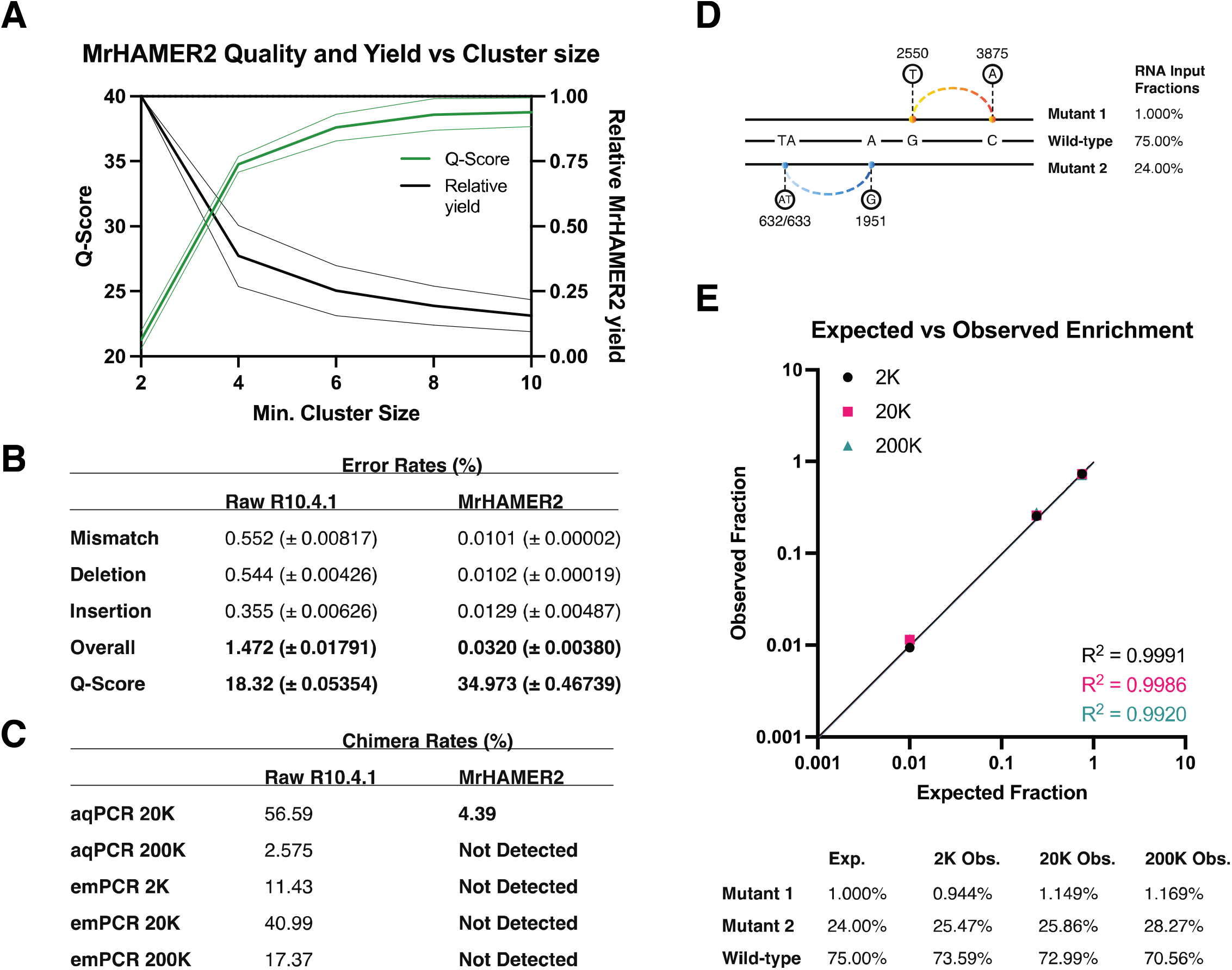
MrHAMER2 yields highly accurate single-molecule long-read RNA sequences across a range of concentrations. **(A)** Cluster size is directly proportional to MrHAMER2 error correction levels in terms of Quality Scores. Despite increased single-molecule accuracy, larger cluster sizes also reduce the yield of MrHAMER2 reads. A cluster size of 4 shows a reasonable balance of accuracy and yield of MrHAMER2 reads. **(B)** Error profiles at a cluster size of 4 show that indels are a dominant error type, constituting around 66% of total errors, consistent with the error profiles observed in Raw R10.4.1 reads. MrHAMER2 reduces the error rate of stock Nanopore chemistries from 1.5% to 0.03%. **(C)** Post-hoc chimera filtering using dual UMIs reduces PCR recombination artefacts to undetectable levels when using emulsion PCR (emPCR) and when using higher RNA input during aqueous PCR (aqPCR). When using aqPCR with lower input amounts, chimera filtering reduces chimeras by 12.9-fold. **(D)** Three mutants containing distant linked single nucleotide variants (SNVs) are admixed at indicated proportions to generate a synthetic RNA reference dataset to test long-range linkage detection and fidelity. **(E)** MrHAMER2 accurately captures long-range linkage across a range of enrichment levels, with sequences detected at expected frequencies (R2 > 0.99) when using 2K, 20K, and 200K RNA molecules. Values in panel (A) are means ± 95 CI. Values in panels (B) and (C) are medians ± SD.

### Validation of isoform quantification accuracy and sensitivity using biologically relevant PCR-free reference data

We next set out to validate that MrHAMER2 can accurately detect, sequence and quantify the full breadth of HIV-1 transcripts which, despite their very low enrichment in cellular mRNA^5, 6^, show prodigious isoform diversity (up 100 isoforms) and dynamic range^7^. Given that no synthetic RNA reference datasets are available for spliced HIV isoforms, we opted to generate a PCR-free reference using our Direct cDNA sequencing protocol, previously validated to yield full-length spliced transcripts in an unbiased and quantitative manner^5^. To ensure that the reference data are biologically relevant and yield the full gamut of HIV-1 isoform diversity, we used primary cells derived from three healthy donors which were then productively infected *ex vivo* with the NL4-3 lab-adapted HIV-1 isolate. To evaluate the potential limits of detection of the MrHAMER2 approach in capturing the gamut of full-length isoforms, the fraction of HIV-1-infected cells from each cell pellet was evaluated using digital droplet PCR (ddPCR) **(Table 1)**, showing that a median of 50,000 HIV-1+ cells were present in the typical cell pellets used for sequencing. Total RNA from these samples was sequenced to saturation with our PCR-free scheme in the P2 Solo and in parallel in the MinION with our MrHAMER2 approach. HIV-1 reads from both datasets were processed and analyzed using the Analysis Pipeline for HIV Isoform eXploration (APHIX)^18^, a fully automated and optimized implementation of our previous HIV isoform quantification workflow^5^. We found that across HIV-1 splice junctions, MrHAMER2 results were very highly correlated with those in the PCR-free reference across 4-logs of enrichment **(Fig. 3A)**, with R^2^ values of 0.9928, and splice junction usage consistent with values previously reported for active infection by us and others for highly enriched junctions (such as D4|A7) and rare junctions (such as D2b|A3)^7, 10, 11^. By using long-reads containing full exon connectivity, isoforms can be unambiguously assigned to likely gene products by identifying the first undisrupted ORF closest to the 5’ end of the transcript^5^. We found that isoform usage in MrHAMER2 reads was highly correlated with those in our PCR-free reference across 3-logs and with R^2^ values of 0.9934 **(Fig. 3B)**. These isoform assignment numbers are also highly concordant with those found by us and others in the context of productive infection^7, 9, 12^, further highlighting the relevance of our validation approach. Overall, these results demonstrate that MrHAMER2 can accurately capture and quantify long-read isoforms with high dynamic range from cellular RNA mixtures obtained from biologically relevant isolates, even when target RNA sequences are only present in a small subset of cells.

**Figure 3.**
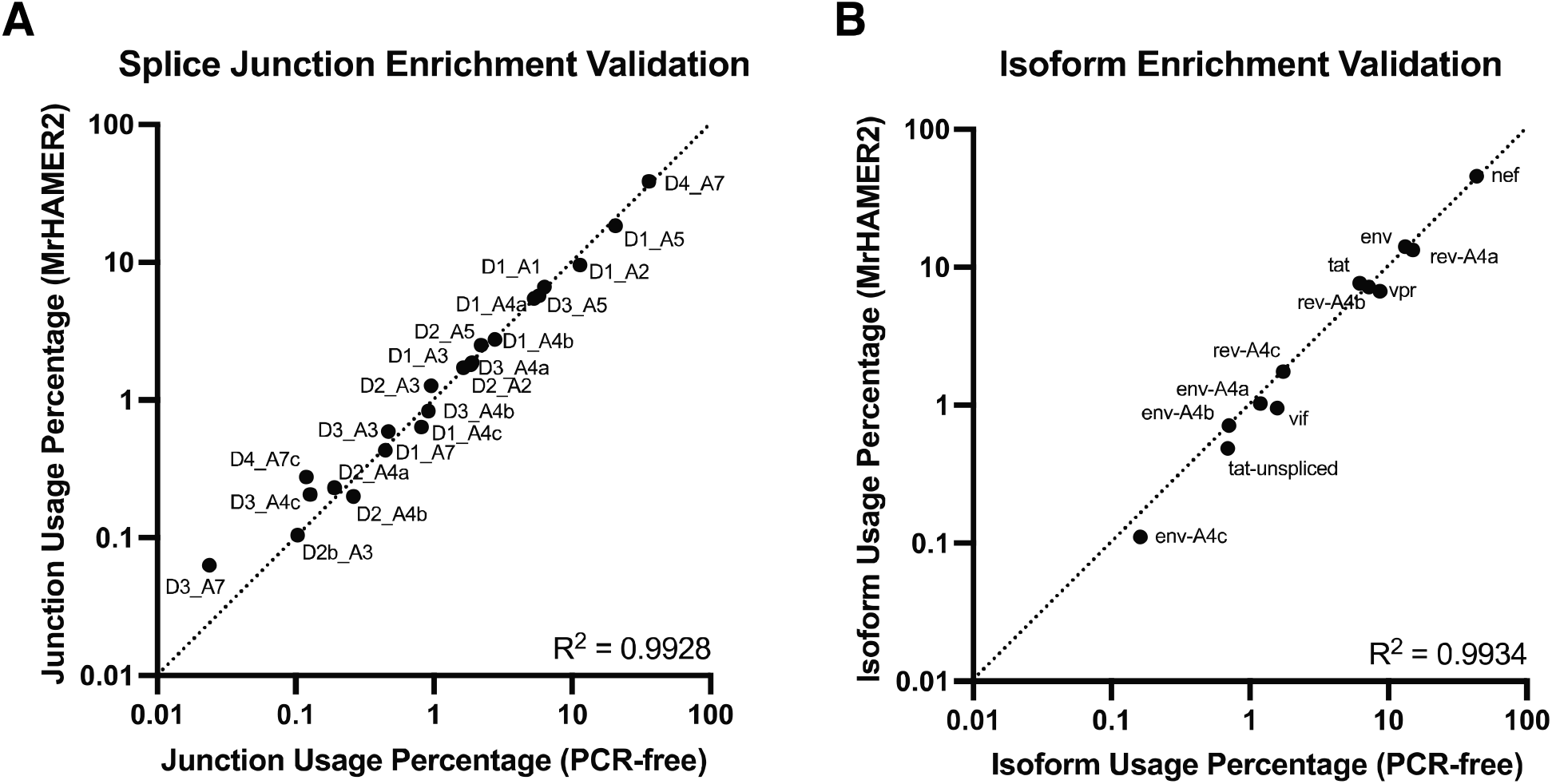
MrHAMER2 can accurately detect and quantify full-length viral isoforms with high dynamic range when benchmarking against matched PCR-free reference data from primary cells. **(A)** Comparison of splice junction usage in a matched biological sample between a PCR-free reference dataset generated using our previously validated approach^5^ and MrHAMER2. Splice junction usage fractions are highly correlated (R^2^>0.99) between MrHAMER2 and the PCR-free reference across the 4-log dynamic range. **(B)** Isoform assignment, as analyzed via our APHIX^18^ pipeline, yields highly correlated isoform enrichment between PCR-free data and MrHAMER2 (R^2^>0.99), with values of most enriched and less enriched isoforms consistent with ranges previously observed by us^5^ and others^7, 10^.

**Table 1.** Cell equivalents and estimated HIV-1+ cells used for MrHAMER2 sequencing of actively infected cells.

| **Sample** | **Cell equivalents** | **HIV DNA per cell** | **Approx. HIV+ cells** |
| --- | --- | --- | --- |
| **Donor1_Productive** | 10,285 | 0.019 | 192 |
| **Donor2_Productive** | 7,571 | 7.009 | 53,061 |
| **Donor3_Productive** | 11,738 | 7.192 | 84,412 |
| Cell equivalents for a given sample are calculated by multiplying the following values: (A) the fraction of total RNA isolate that was used for Reverse Transcription, (B) the subsequent fraction of dual UMI-tagged cDNA that was used to generate a well-sampled MrHAMER2 library, and (C) the cell number of the pellet from which total RNA was isolated. The number of HIV+ cells was obtained by multiplying the HIV DNA per cell (as determined via digital droplet PCR) by the Cell equivalents. |  |  |  |

### MrHAMER2 decodes the HIV-1 transcriptional program in a primary CD4+ T cell derived viral latency model

Having validated that MrHAMER2 can accurately quantify isoform-specific variation in viral transcripts from biologically relevant samples, we set out to interrogate the spliced HIV-1 transcriptome in the context of viral latency. We used a well-characterized model of latency^19^ where primary CD4+ T cells are *ex vivo* infected with HIV-1, activated with anti-CD3 and anti-CD28 antibodies, and mixed with autologous resting uninfected cells in a 1:4 ratio **(Supplementary Fig. 4)**. This admixing results in cell-to-cell infection of resting cells and generation of a clear viral latency phenotype in this cell fraction^20^. Using ddPCR we found that bulk latently-infected cells show up to 400-fold lower fraction of HIV-1+ cells compared to active infections **(Table 2)**, which translates to as low as 58 HIV-1+ cells per typical cell pellet used for MrHAMER2 sequencing.

**Table 2.**
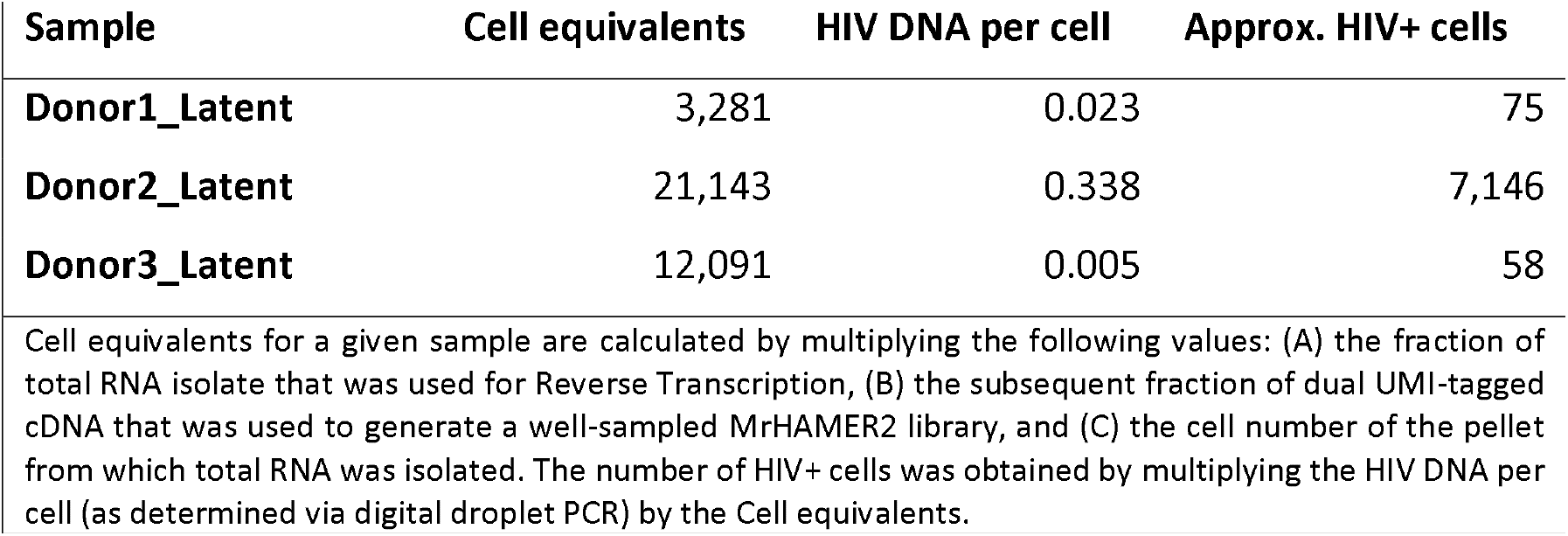
Cell equivalents and estimated HIV-1+ cells used for MrHAMER2 sequencing of latently-infected cells.

Our model of viral latency is characterized by low levels of HIV transcriptional activity, which is beneficial for the evaluation of HIV-1 isoform dynamics during latency^19, 20^. We first used MrHAMER2 to compare HIV-1 isoform dynamics in productively infected CD4+ T cells to autologous latently-infected CD4+ T cells. We determined that the viral isoform signature in latently-infected CD4+ cells **(Fig. 4A)** was characterized by statistically significant (at least p<0.01) increases in Env, Tat, and Vif isoforms (34%, 128%, and 488% increases respectively), along with a concomitant decrease in Nef, and Rev-A4a isoforms (28% and 40% decrease respectively). Due to the high accuracy of MrHAMER2 reads, isoform productivity (i.e. whether a transcript has the potential to generate a translatable protein) can be determined at the single-molecule level **(Fig. 4B)**. Surprisingly, we found that MrHAMER2 reads belonging to Env clusters have significantly lower isoform productivity of <60% compared to other isoforms which average greater than 90% productivity. More importantly we found isoform productivity differentials between productively and latently-infected cells, with Env and Vif isoforms showing significantly lower isoform productivity by 15% in productive compared to latent infection (p<0.001 for both isoforms). To trace the potential causes of these differences in isoform productivity, we performed an error profile analysis of MrHAMER2 reads belonging to Env and Nef isoform clusters, which have identical exon structures except for a single IR event (D4|A7) in the former. Error profile analysis revealed that the differences in isoform productivity may be explained by a doubling in median mutational burden in Env isoforms compared to Nef isoforms **(Supplementary Fig. 5)**.

**Figure 4.**
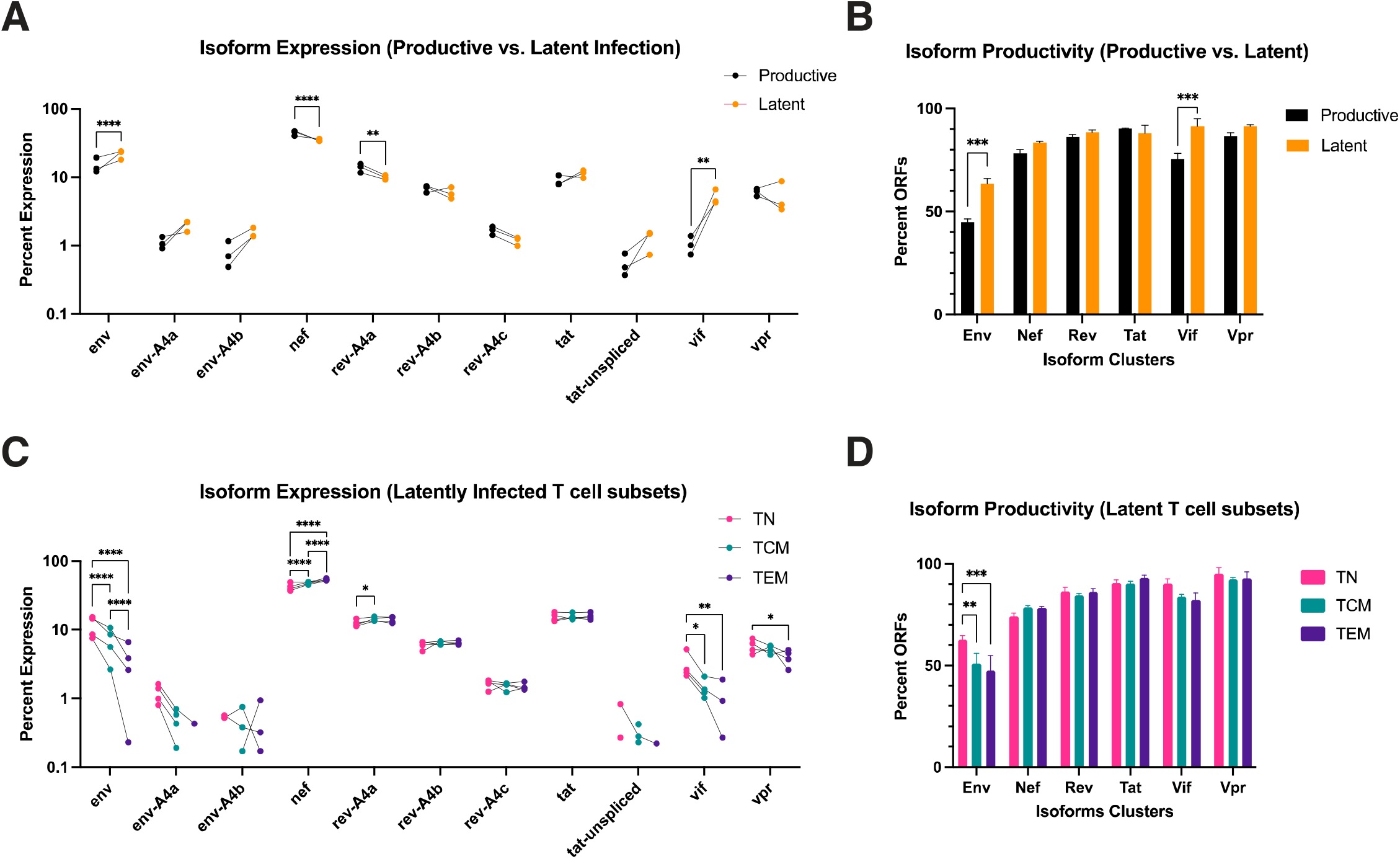
MrHAMER2 decodes the viral transcriptional program in primary CD4+ cell derived model of HIV-1 latency. **(A)** Isoform expression comparison between matched autologous samples in productive and latent infection. Statistically significant changes in Env, Nef, Rev, and Vif isoforms are observed, with the largest magnitude changes observed in Env and Vif isoforms, which contain intron retentions. **(B)** Isoform Productivity (i.e. the translatability of isoforms) was calculated by computing the percent of isoforms that can code for Open Reading Frames (ORFs) as a fraction of the total isoforms within an isoform cluster. Significant reductions in isoform productivity are observed in Env and Vif isoforms in productive compared to latent infection. **(C)** Isoform expression signatures in latent T cell subsets shows order of magnitude changes in isoforms containing IRs, including Env, Vif and Vpr. These changes in IR-containing isoforms results in a concomitant Nef isoform expression differential (TN, naïve; TCM, central memory; TEM, effector memory cells). **(D)** Isoform Productivity in latently-infected T cell subsets recapitulates the lower percent ORFs found in Env isoforms in productively infected and unsorted latently-infected samples. A slight but statistically significant isoform productivity differential was observed between TCM and TEM subsets compared to TN. Data in panels (A) and (B) are from three biological replicates. Data in panels (C) and (D) are from four biological replicates. Values in panels (B) and (D) are means ± SEM. Statistical significance was calculated with two-way ANOVA with Tukey multiple comparison test: *P< 0.05; **P< 0.01; ***P< 0.001; ****P< 0.0001.

Despite the utility of latently-infected primary CD4+ T cells in identifying general transcriptional regulatory trends involved in latency, the viral isoform signatures identified in these samples can be heavily biased by the CD4+ T cell compositions of each donor. To evaluate the potential interplay between CD4+ T cell effector/memory functions and viral isoform signatures, latently-infected cells from each donor were sorted into Naive (TN), Central Memory (TCM), and Effector Memory (TEM) subsets and then sequenced via MrHAMER2. As expected, TN cells were identified as a major fraction of latently-infected cells with up to 50% enrichment **(Supplementary Fig. 6)**. Importantly, TCM cells, a putative viral reservoir^21^, were found less enriched at ∼25%, while TEM cells were a minority fraction at ∼4%. Despite the broad differences in enrichment levels in these subsets, all canonical viral isoforms were detected and quantified across samples from 4 donors. We found that viral isoform signatures in TCM and TEM subsets were characterized by an order of magnitude decrease of all partially spliced transcripts (Env, Vpr, Vif) compared to their respective TN subsets (p<0.0001 in Env, p<0.01 in Vif and Vpr), suggesting that viral gene expression in these subsets may be modulated by changes in IR **(Fig. 4C)**. The decrease in isoforms that harbor IRs, was accompanied by a concomitant increase in Nef isoform of around 60% (p<0.001). The lower Env isoform productivity previously observed in productively infected and unsorted latently-infected cells are recapitulated by CD4+ T cell subset data with an average 50% productivity observed in all subsets, and modest but statistically significant (at least p<0.01) reductions in productivity in Env in TCM and TEM compared to TN counterparts **(Fig. 4D)**. The substantial isoform differentials gleaned from MrHAMER2 sequencing, particularly in viral isoforms involved in antigen presentation^22^ and cell-cycle arrest^23^, suggests that an immune evasion phenotype in TCM or TEM subsets could be a driver of HIV-1 persistence in primary cells.

## DISCUSSION

We developed MrHAMER2, a highly accurate (Q35) long-read sequencing pipeline, to detect and quantify isoform-specific variation in viral transcripts in contexts of low transcriptional activity. The pipeline builds on our previous long-read RNA sequencing developments^5, 6, 14, 24^, and couples them with dual UMI tagging of single cDNA molecules. MrHAMER2 enables the accurate amplification of the full breadth of viral isoforms from low abundance RNA while controlling for PCR sampling bias^25^ and recombination artefacts. Applying MrHAMER2 to a primary CD4+ T cell-derived model of viral latency^20^, we provide, to our knowledge, the first comprehensive characterization of full-length viral transcriptional program and associated isoform productivities during HIV-1 latency in biologically relevant CD4+ T cell subsets, a knowledge gap that had been unaddressed due to technical limitations.

Using MrHAMER2 in a primary CD4+ T cell model of HIV-1 latency, we believe we are the first to describe that substantial changes in viral isoforms containing IRs underlie the transition to viral quiescence in primary CD4+ T cells. Given the role of cellular latency in the maintenance of the viral reservoir, understanding associated viral isoform signatures could provide clues on the post-transcriptional regulation processes that underlie viral persistence. IRs have been previously shown as a mode of post-transcriptional regulation during active infection in both host-cell^22^ and viral^26^ transcripts. However, the role of IR in latency has only been studied indirectly in the context of treatment of HIV-1-infected cells with Filgotinib^27^ (a JAK inhibitor) and Topotecan^28^ (a Camptothecin analog) which have been proposed as HIV-1 suppressing agents. Both Filgotinib and Topotecan selectively decrease fully spliced viral RNA, indirectly enriching for less processed viral RNA containing IRs. Our data from latently-infected CD4+ T cells **(Fig. 4A)**, showing increases in viral isoforms containing intron retentions (Env, Vif), along with decreases in fully spliced species (Nef, Rev-A4a) are consistent with the model that Filgotinib and Topotecan further promote HIV-1 latency via mechanisms such as IR. Our data in CD4+ T cell TCM and TEM T cell subsets **(Fig. 4C)**, showing almost 0.5-1.0-log reductions in IR-containing isoforms (Env, Vif, Vpr) are consistent with the results reported using Ruxolitinib, another JAK inhibitor, in the context of HIV-1 suppression^27^. Our findings are suggestive of a mechanistic role of intron retention as a possible explanation for the dynamics of associated viral gene products and their downstream functional effects in immune activation^29, 30^, immune evasion, or replication capacity. The findings also indicate that intron retention may be a possible biomarker of viral latency.

To account for future improvements in Nanopore sequencing accuracies, the bioinformatic components in MrHAMER2 are modular so that polishing models associated with basecalling updates can be downloaded and invoked in the pipeline’s configuration file. For example, recent updates in Oxford Nanopore Technologies (ONT) sequencing chemistries and basecalling implementations have reduced raw single molecule errors to 1.5%^31^. These improvements allowed us to attain better MrHAMER2 single nucleotide accuracy (Q35 vs Q30) and higher read yield (10-fold higher) with smaller cluster sizes per single molecule (4 vs 10) compared to the first generation of this protocol^14^. Despite these accuracy improvements, indels remain the dominant error mode at ∼66% of all errors. These error profiles might be improved by rapid developments in ONT sequencing chemistries^32^ or basecalling implementations^33^ (including species- or strain-specific basecalling models^34, 35^). Additional Nanopore platform accuracy improvements will translate to better single nucleotide accuracy when using MrHAMER2 or, alternatively, allow for smaller cluster sizes to obtain a higher yield of MrHAMER2 reads. We have seen an early display of such improvements in yield vs. accuracy when using a dual UMI-assisted duplex basecalling^36^ to “rescue” single molecule clusters of size two (that otherwise would have been filtered out) and observe at least doubling in MrHAMER2 read yield while retaining reasonable Q33 accuracy **(Supplementary Fig. 3b)**.

Future studies could leverage MrHAMER2’s high sensitivity to obtain both cellular and viral transcriptional landscapes at the single-cell level. Given the sensitivity of MrHAMER2 to obtain the full breadth of viral isoforms from as few as 58 HIV-1 positive cells, coupling the approach with single-cell RNA sequencing should be tractable^37, 38^. This would provide insights into specific cells and subsets that have differential viral RNA burst sizes and determine associated host-cell transcriptional signatures that sustain viral replication or latency. On the other hand, studies on RNA structure determinants of splicing via isoform-specific RNA structure probing could also be attempted by taking advantage of the substantial improvements in signal-to-noise ratio afforded by the high single-nucleotide accuracies in MrHAMER2.

## DATA AVAILABILITY

Sequencing data have been submitted to the European Nucleotide Archive (ENA) under study accession PRJEB88729. Both raw nanopore data in FAST5 format and basecalled data in FASTQ format are available. MrHAMER2 bioinformatic pipeline is available at www.github.com/gallardo-seq/MrHAMER2. APHIX bioinformatic pipeline for viral isoform analysis is available at www.github.com/jessicaA2019/APHIX.

## ACKNOWLEDGEMENTS

We thank Dr. Maile Karris, Ms. Deedee Pacheco and the San Diego CFAR Clinical Investigation Core for recruiting study participants and providing peripheral blood samples. We thank the San Diego CFAR Molecular and Cellular Immunology Core for support in conducting cell sorting and ddPCR experiments.

## MATERIALS AND METHODS

### Primers and oligos

GSP_UMI_Fwd_2.U5.B4F (PAGE Purification): GTATCGTGTAGAGACTGCGTAGGTTTVVVVTTVVVVTTVVVVTTVVVVTTTAGTAGTGTGTGCCCGTCTGTTGTGTGACTC

GSP_UMI_Rev_3’LTR (PAGE Purification): AGTGATCGAGTCAGTGCGAGTGTTTVVVVTTVVVVTTVVVVTTVVVVTTTTAACCAGAGAGACCCAGTACA

UVP_Fwd (Standard Desalting): GTATCGTGTAGAGACTGCGTAGG UVP_Rev (Standard Desalting): AGTGATCGAGTCAGTGCGAGTG

### Total RNA isolation

Total RNA was isolated from cell pellets using the RNeasy Mini Kit (QIAGEN, cat #74134). Cells were lysed with RLT buffer (with no β-ME), processed according to manufacturer’s instructions, and eluted in 50 μl nuclease-free water. Total RNA sample quality was assessed via gel electrophoresis with E-gel EX 1% system and shown to have a 28S to 18S rRNA ratio greater than or equal to 2.7 for all samples.

### Chemical ablation of 3’ RNA ends (CASPR)

Sodium periodate (NaIO4) was purchased from Millipore Sigma (311448-5G). A 2× buffered periodate solution (BP) was prepared fresh each time by measuring NaIO4 powder and resuspending to a concentration of 4 mg/ml in aqueous solution of 200 mM sodium acetate (pH 5.5) (Invitrogen, AM9740). Input RNA (up to 5 μg) was mixed with an equal volume of 2× BP and incubated at room temperature in the dark for 30 min. Following treatment, RNA was cleaned with RNA Clean & Concentrator (Zymo Research, R1013) according to the manufacturer’s instructions, and eluted in nuclease-free water.

### Reverse transcription

Reverse transcription was carried out with SuperScript IV Reverse Transcriptase (Invitrogen 18090050) in a 20 μl volume with the following components and final concentrations: 1× Reaction Buffer, dNTPs (0.5 mM), RNAseOUT (2U/μl), Oligo-d(T)_20_ (0.5 μM), CASPR-treated total RNA (≤500 ng), DTT (5 mM), and SuperScript IV RT (10 U/μl). Primer was initially annealed to template RNA in the presence of dNTPs, by heating to 65°C for 5 min, followed by snap cooling to 4°C for 2 min. After snap cooling, the rest of the components were added, followed by reverse transcription for 1.5 h at 50°C. Reactions were stopped by heat inactivation at 85°C for 5 min. Reactions were cleaned with Monarch DNA Clean kit and eluted in 10 μl EB. First-strand products were cleaned with 1X AMPure XP beads according to the manufacturer’s instructions, then eluted in 40 µL of 0.1X TE.

### Dual UMI tagging

Dual UMI tagging was carried out with Phusion U Hot Start Polymerase in a 100 µL volume with the following components and final concentrations: 1X GC Buffer, dNTPs (0.2 mM), GSP_UMI_Fwd_2.U5.B4F (0.1875 µM), GSP_UMI_Rev_3’LTR (0.1875 µM), RNAse H (5 U), RNAse If (50 µL), Phusion U Hot Start DNA Polymerase (2 U). Samples were thermally-cycled through following program: 37°C (15 min), 98°C (1 min), 72°C (10 min), 98°C (1 min), 63°C (30 sec), 72°C (10 min), 4°C Hold. After cycling 5 µL of Themolabile Exo I was added to each sample and then incubated at 37°C for 5 mins, followed by heat inactivation at 80°C for 2 mins. Dual UMI-tagged samples were then cleaned with 1X AMPure XP cleanup and eluted in 40 µL 0.1X TE.

### Emulsion PCR

Aqueous phase was prepared in a 1.5 ml DNA LoBind tube to a final volume of 50 μl with the following components and final concentrations: cDNA input, 1X Phusion GC Buffer, 0.2 mM dNTPs, 0.5 μM each of UVP_Fwd and UVP_Rev primers, 0.5 mg/ml BSA, and 0.02U/μl Phusion U Hot Start DNA Polymerase (F555S). Oil/Surfactant was prepared: 2% (v/v) ABIL em90 (Evonik Degussa GmbH) and 0.05% (v/v) Triton X-100 in Mineral oil (31). Three hundred microliters of Oil/Surfactant were added on top of the aqueous phase, then vortexed for 5 mins at maximum speed using Vortex-Genie 2. Emulsified components were aliquoted into PCR strip (each tube containing no more than 50 μl). Samples were thermally cycled through following program: 98°C (2 min), 98°C (10 sec)/62°C(30 sec)/72°C (7:30 min) for 25 cycles, 72°C (5 min) and 4°C Hold. Following thermal cycling, emulsion was consolidated in a 2 ml DNA LoBind tube and broken by adding 700 μl ethyl acetate and vortexing for 5–10 s. One milliliter of DNA binding buffer (Monarch DNA Clean) was added followed by vortexing for 10 s. Tube was spun at 20 000 x g for 2 min, resulting in three phases. Aqueous phase, which settled in bottom of tube, was carefully aspirated, and transferred directly to the DNA clean column with cleanup washes proceeding as indicated on manufacturer’s protocol, followed by elution in 0.1X TE buffer. Eluted DNA from emulsion was further cleaned with 0.66X AMPure XP beads and eluted in 22 µL 0.1X TE. 2X cleaned DNA was ready for evaluation using agarose gel electrophoresis with E-Gel EX 1% gel system and via QuBit dsDNA HS. If yield was less than 50 ng, then a second 10-cycle emulsion PCR was performed using 1-5 ng of input.

### Nanopore sequencing and basecalling

All MrHAMER2 samples were barcoded and library prepped with Native Barcoding Kit 24 V14 (SQK-NBD114.24). All samples were sequenced with MinION R10.4.1 flow cells, basecalled and demultiplexed with Guppy or Dorado basecallers using “sup” basecalling models.

### Bioinformatic processing

Basecalled reads were concatenated into a combined FASTQ file for each sample. Data were then processed using the MrHAMER2 snakemake pipeline (https://github.com/gallardo-seq/MrHAMER2) by providing input FASTQ files for each sample, the NL4-3 Reference sequence, and invoking both of these files in the Snakemake configuration along with parameters such as “min_reads_per_cluster” and “balance_strands=True”. A min_reads_per_cluster=2 was used for counting applications (such as isoform enrichment) and a min_reads_per_cluster=4 was used for single-nucleotide variation analyses (such as isoform productivity). Resulting high accuracy MrHAMER2 reads were then processed with the APHIX^18^ pipeline (https://github.com/JessicaA2019/APHIX) using an APHIX cluster_size=2 to obtain quantification and analysis of isoform and splice junction usage, and isoform identity and enrichment. Detailed installation and running instructions for MrHAMER2 and APHIX bioinformatic pipelines are provided in their respective GitHub links.

### Dual UMI-assisted duplex basecalling

MrHAMER2 bioinformatic pipeline was run with a min_reads_per_cluster=2 and “balance_strands=True” parameters. The resulting MrHAMER2 read IDs from FASTQ output were then cross-referenced with the “id_cluster” column in “*runID*_vsearch_cluster_stats.tsv” file in the /stats/ subdirectory, and a list of id_cluster with a value of 2 in the “written column” was exported into a separate txt file. The resulting cluster_ids.txt file, containing a list of clusters that have exactly one sense and one antisense read, was then used to extract the read_IDs from the fasta files that were generated for each id_cluster in the /clustering/*run_ID*/clusters_fa/ subdirectory during MrHAMER2 processing, and a script was used to list sense and antisense read ID pairs in a txt file per line (with each line corresponding to sense and antisense read_IDs for a given id_cluster). The resulting pairs text file was then used to invoke duplex basecalling with guppy_basecaller_duplex (version 6.1.7) using the following arguments: -i fast5_directory -s output_directory -c basecalling_model.cfg --recursive --duplex_pairing_mode from_pair_list --duplex_pairing_file pairs.txt.

### Isoform productivity analysis

MrHAMER2 reads obtained with a min_reads_per_cluster=4 (for high accuracy SNV analysis) were processed via APHIX pipeline (with an APHIX cluster_size=2) to yield isoforms belonging to specific viral genes (Env, Nef, Rev, Tat, Vpr, Vif). The MrHAMER2 reads belonging to each isoform were then extracted and respectively analyzed for their ability to generate intact ORFs using the ‘ORF Finder’ tool in Geneious, with the following filtering parameters per isoform/gene product: Env (2500 bp min length), Nef (600 bp min length), Rev (275-425 bp start site, 630-800 bp end site, 348-375 length), Tat (261 bp min length), Vpr (275-300 bp length, min length less than 610 bp), Vif (length 570-600 bp). The ORF Finder filtering criteria for each isoform cluster was validated by cross-referencing the resulting protein sequences with those reported in UniProt. Isoform productivity was then computed by taking a ratio of the number of reads that result in translatable product to the total number of MrHAMER2 reads within an isoform cluster.

### Generation of latently-infected cells

Cells were cultured in RPMI + 5% human AB (HAB) serum, which has pen/step/glutamine supplement added to it in all cases. On Day -1, CD4+ T cells were isolated from blood (300 ml) of HIV seronegative donors by negative selection (StemCell Technologies, Inc) yielding ∼100 million cells. Purity and activation were verified with a panel of antibodies (CD4+-FITC, CD8-PE, CD19-FITC, CD16-PE, CD3-APC, HLA-DR-PE) and samples proceeded to the next step only if purity was >90% CD4+ T cells and activation was <10% HLA-DR+. 20 million cells were then stained with the eFlour 670 viability dye. Remaining cells were resuspended in conditioned media (4 volumes of RPMI + 5% HAB + 1 volume of conditioned media) at 5 million cells per 1.5 ml, and incubated in 12 ml tubes with 1.5 ml per tube at 37°C until Day 4. On Day 0, eFluor-670 stained cells and a small aliquot (5×10E6) unstained cells were infected with NL4-3 virus and incubated at 37°C for 4-6 hrs. Cells were then washed 4 times with PBS+2%HAB to remove any virus that did not enter cells. During infection, plates were prepared for cell activation by coating with goat anti-mouse antibody (>90 min), washing with PBS, blocking with PBS+2%HAB (>30 min), washing with PBS, then incubating with anti-CD3/anti-CD28 antibodies (1:200 and 1:500 dilutions) for at least 1 hour at room temperature and washing with PBS right before plating. Cells were washed 2x with PBS then plated in the following manner: uninfected (control), infected unstained (control for flow sort), and infected stained cells. On Day 4 resting cells were pooled in conditioned media into a single tube, while stained/infected/activated cells were consolidated into a separate tube. Resting and activated cells were counted, then mixed at a 4:1 ratio (while saving some resting cells for flow sort control). Samples from all conditions were taken for p24 staining to evaluate levels of infection. Mixed cells were spun down and resuspended in RPMI + 5%HAB + IL2/IL15 at 5 million cells per 3 ml media. Cells were plated in 6-well plates at 3 ml per well. On Day 7, cells were collected from plates and submitted to the Molecular and Cellular Immunology Core to sort unstained cells, which were never activated, but acquired virus via cell-to-cell transmission from productively infected cells. After the sort, unstained cells were resuspended in RPMI+5% HAB and incubated in tubes (1.5 ml per tube, 5 million cells per tube) until day 10. A small aliquot of cells from all samples (including controls) were saved for assessment of p24 production. On Day 10, HIV-1 latently-infected cells were collected for downstream assays. A small aliquot (500,000 cells) was collected for integrant DNA assay to assess the reservoir size, and the rest of the cells were lysed with RLT Buffer (without ß-ME) for long transcript sequencing.

### Generation of productively infected cells

Extra unstained infected cells were activated (see latently-infected section above), so they could also be collected at Day 7 for long transcript sequencing. Briefly, on Day -1 CD4+ T cells were isolated from blood (300 ml) of HIV-1 seronegative donors by negative selection (StemCell Technologies, Inc). On Day 0, 5 million unstained cells that were infected and activated were set aside. On Day 4, infected cells were collected from plate and resuspended in RPMI + 5% HAB + IL2/IL15. On Day 7, these cells were collected for long transcript sequencing. 500,000 cells were set aside at this point for integrant DNA assay.

### Sorting of T cell subsets

In experiments with CD4+ T cell maturation subsets, TN, TCM and TEM cells were sorted on day 10 using flow cytometry. Before the sort, an aliquot of cells was taken for long transcript sequencing of the total cell population. The remaining cells were stained with CD4+5RA-PE-Cy7 and CD27-APC antibodies and 3-way sorted into TN (CD4+5RA+ CD27-APC+), TCM (CD4+5RA-CD27-APC+) and TEM (CD4+5RA-CD27-APC-) populations.

### Quantification of HIV integration via ddPCR

Total genomic DNA was extracted from cell pellets using Qiagen QIAamp DNA Micro Kit and the high molecular weight fraction was purified using a short-read elimination kit (PacBio), which depletes the sample from all DNA fragments <10kb. ddPCR was used to quantify copies of HIV DNA (GAG and 2LTR circles) in each sample. Ribonuclease P/MRP subunit P30 (RPP30) host genomic DNA was quantified to estimate the total number of cells contributing to each reaction. Integrated HIV DNA copy numbers were corrected by subtracting the 2LTR circles and normalized to the cellular DNA input.

