## Supplementary Figures for "MrHAMER2: high-accuracy long-read RNA sequencing to decode isoform-specific variation in viral transcripts during latency"

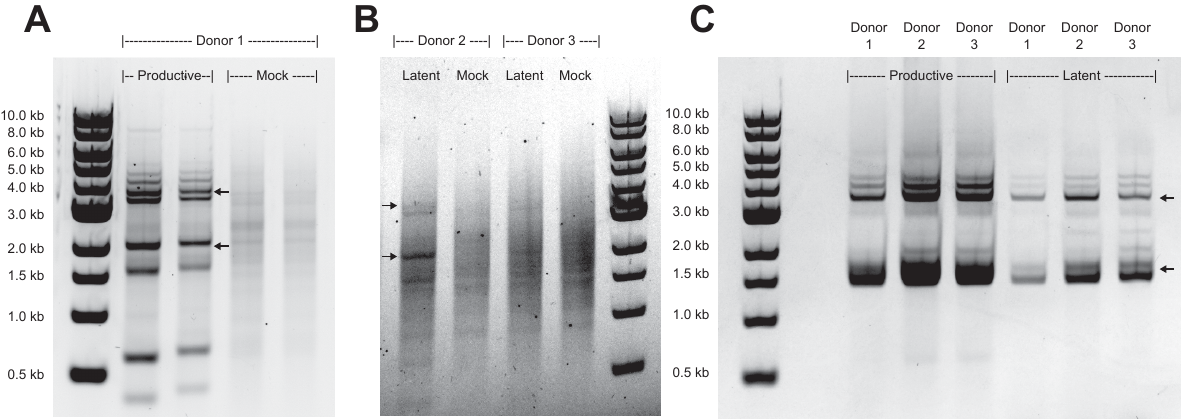

**Supplementary Figure 1 - MrHAMER2 priming strategies tested in productively and HIV-1 latently infected cells. (A)** 3' UMI is added during Reverse Transcription using Oligo-d(T) priming, followed by 5' UMI tagging via gene-specific priming during second strand synthesis. This priming strategy results in emPCR amplicons with major two isoform clusters (see arrows) of expected size and of sufficient signal when using productively infected samples. However, significant background is observed in mock infected samples despite the lack of viral RNA present in these cells. **(B)** Gel of PCR products using the UMI tagging and priming strategy in (A) when using Latently infected cell samples from two donors. In this case, very low signal is obtained from Donor 2, while insufficient signal is obtained from Donor 3. Both samples have significant amount of background signal that corresponds with that observed in respective mock samples. This suggests this priming strategy is not suitable with samples having low transcriptional activity. **(C)** Improved UMI tagging strategy involves conventional Oligo-d(T) primed RT without flanking adapters, followed by sequential addition of 5' and 3' UMI tags during second strand synthesis. Gel products show two isoform clusters of expected size (see arrows) in both productively and latently infected samples without any discernable background.

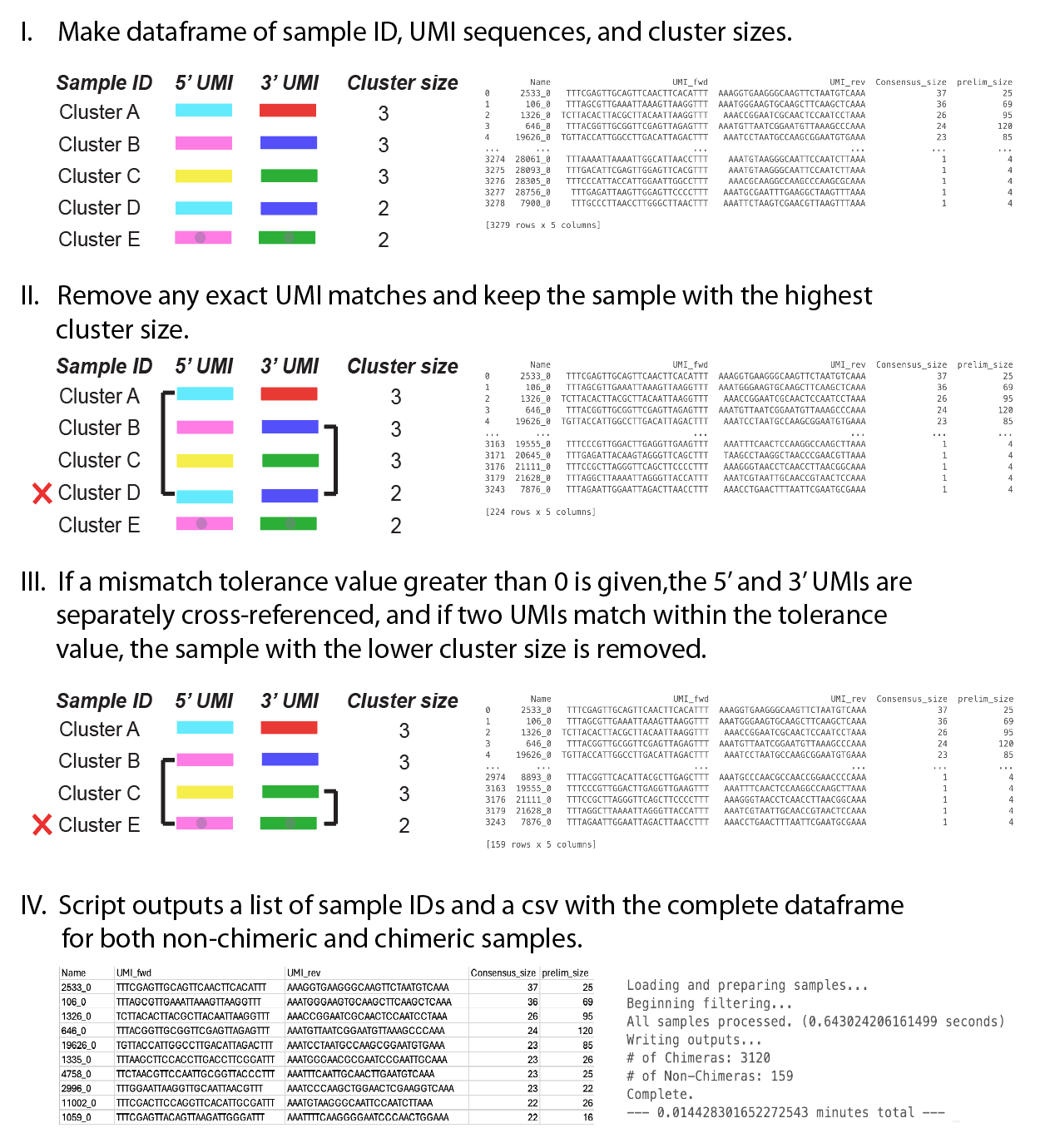

**Supplementary Figure 2 - Schematic of the Chimera Buster module in the MrHAMER2 bioinformatic pipeline.**

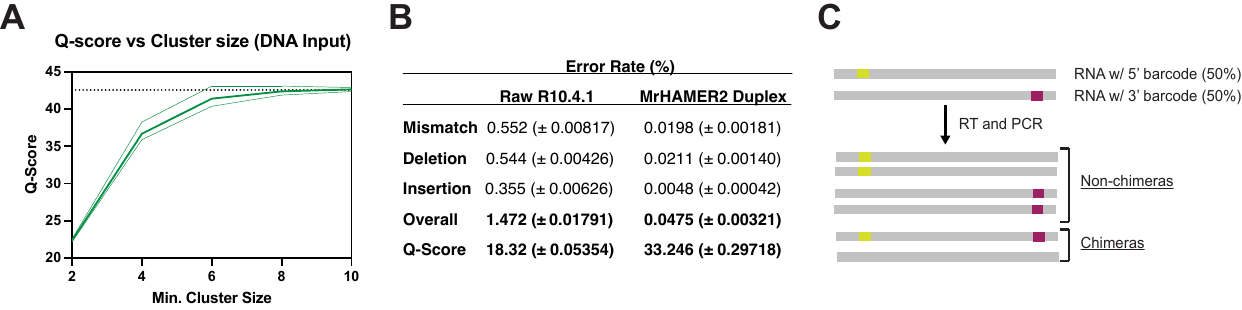

**Supplementary Figure 3 - Additional MrHAMER2 accuracy information and benchmarking. (A)** MrHAMER2 accuracy when using DNA inputs tapers off at around Q42.5. **(B)** Dual UMI assisted duplex basecalling with Guppy basecaller duplex results in Q33 accuracy. **(C)** Synthetic RNA reference dataset used to test chimera filtering capabilities in MrHAMER2.

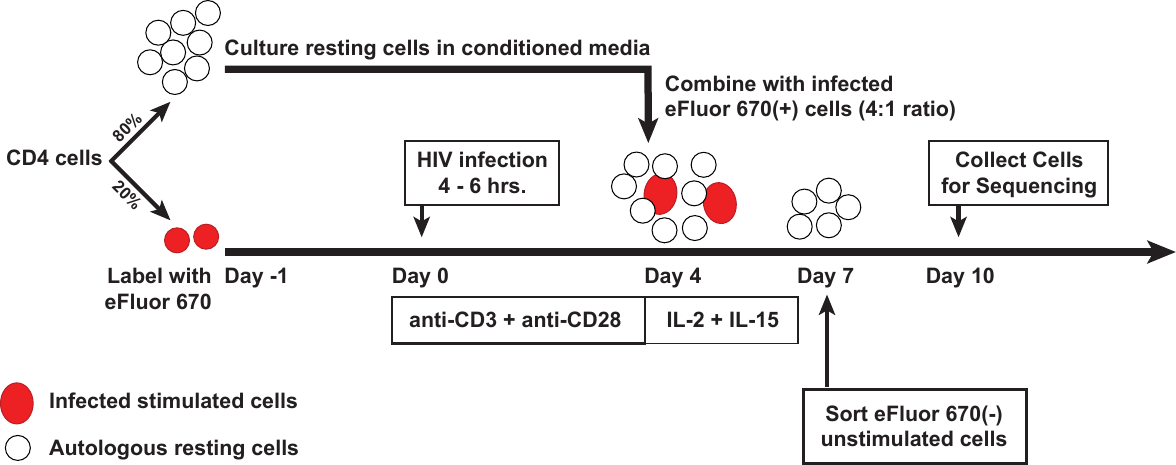

**Supplementary Figure 4 - Schematic of model of CD4+ cell HIV-1 latency used in this study.** Briefly, CD4+ T cells are isolated from healthy donors via negative selection. At Day -1, 80% of the purified CD4 T cells are cultured in conditioned media, while the remaining 20% of cells are labeled with eFluour 670, a cell division marker. At Day 0, eFluor 670 labeled cells are infected with NL4-3 virus for 4-6 hours, and cells are then washed with PBS + 2% human serum to remove any virus that did not enter cells. Infected cells are then incubated in anti-CD3 and anti-CD28 antibody coated plates. At Day 4, resting cells are admixed with infected, activated, eFluor 670-positive cells at a 4:1 ratio. At Day 7, small unstained cells are sorted using flow cytometry, these cells were never activated but were infected via cell-to-cell transmission from the autologous productively infected cells. After the sort, the cells are incubated in media for 3 days. At Day 10 latently infected cells are collected and processed for long-read RNA sequencing via MrHAMER2.

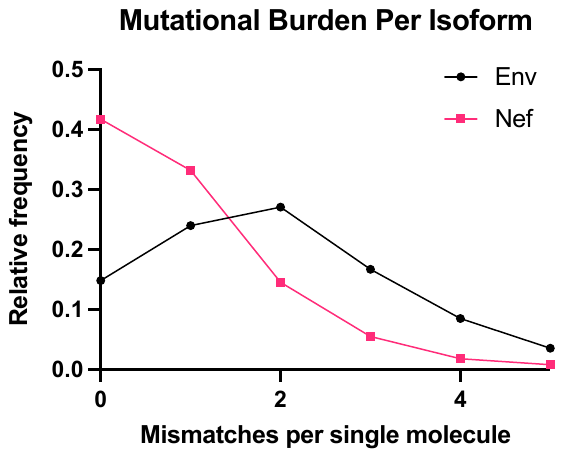

**Supplementary Figure 5. Env isoforms have significantly larger mutational burden compared to Nef isoforms.** Single molecules belonging to Env and Nef isoform clusters are analyzed for number of mismatches per single molecule and a histogram is generated. The majority of Env isoforms have at least 2 mismatches, while the majority of Nef isoforms have less than 1 mismatch, resulting in ≥2X higher mutational burden in the former group.

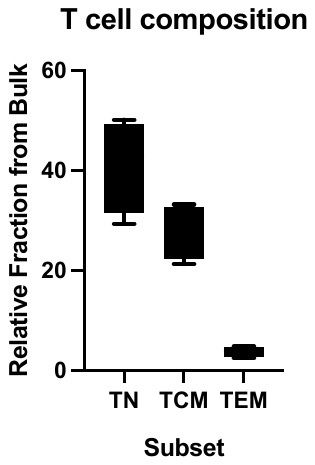

**Supplementary Figure 6.** Approximate CD4+ T cell subsets fractions, Naive (TN), Central Memory (TCM), and Effector Memory (TEM), collected by flow cytometry sorting.
